# Ignoring stratigraphic age uncertainty leads to erroneous estimates of species divergence times under the fossilized birth-death process

**DOI:** 10.1101/477133

**Authors:** Joëlle Barido-Sottani, Gabriel Aguirre-Fernández, Melanie Hopkins, Tanja Stadler, Rachel Warnock

## Abstract

Fossil information is essential for estimating species divergence times, and can be integrated into Bayesian phylogenetic inference using the fossilized birth-death (FBD) process. An important aspect of palaeontological data is the uncertainty surrounding specimen ages, which can be handled in different ways during inference. The most common approach is to fix fossil ages to a point estimate within the known age interval. Alternatively, age uncertainty can be incorporated by using priors, and fossil ages are then directly sampled as part of the inference. This study presents a comparison of alternative approaches for handling fossil age uncertainty in analysis using the FBD process. Based on simulations, we find that fixing fossil ages to the midpoint or a random point drawn from within the stratigraphic age range leads to biases in divergence time estimates, while sampling fossil ages leads to estimates that are similar to inferences that employ the correct ages of fossils. Second, we show a comparison using an empirical dataset of extant and fossil cetaceans, which confirms that different methods of handling fossil age uncertainty lead to large differences in estimated node ages. Stratigraphic age uncertainty should thus not be ignored in divergence time estimation and instead should be incorporated explicitly.

## 1 Introduction

The fossil record provides essential evidence for calibrating species trees to time, as molecular sequences from extant species are informative about the relative age of species but do not typically provide information about the absolute age. A common approach to calibration, referred to as node dating, is to assign a single fossil to a specific node in a phylogeny and to reflect the uncertainty in its age using a probability distribution, where the minimum bound of the distribution corresponds to the age of the specimen (1; 2; 3). It has been shown that divergence time estimates are extremely sensitive to the choice of fossil(s), the age assigned to fossil specimens, and the distribution chosen to model uncertainty (4; 5; 6; 7; 8). However, regardless of specimen choice, node dating has additional drawbacks. For instance, this approach effectively uses one fossil per node and makes it extremely challenging to derive or implement explicit priors on divergence times (9; 10). The fossilized birth-death (FBD) process offers an alternative approach to calibration, which integrates fossil samples into the tree under the same diversification process that describes extant species (11; 12). This approach greatly increases the amount of fossil evidence that can be used during inference, but the impact of the taxonomic, stratigraphic range, and stratigraphic age uncertainty has not been fully explored using this framework.

Here, we explore the impact of stratigraphic age uncertainty. Fossils are rarely composed of material that can be directly dated and their age must be established with detailed reference to the geological record. This procedure leads to some uncertainty. First, the rock layer, or lithostratigraphic unit, from which a specimen was collected must be established. If layers directly above and below that unit have not been directly dated, the relative age, or biostratigraphic unit, of a specimen must be established using index fossils. Finally, the absolute minimum and maximum age of a specimen must be obtained with reference to a numeric timescale, or chronostratigraphic chart. The process of dating fossils is not always straightforward, because the link between litho-, bio- and chronostratigraphy can be challenging to establish, or the stratigraphic provenance of a specimen may be ambiguous (13; 14; 15; 16; 17).

Current applications of the FBD model typically assign specimen ages using the midpoint of the known interval of age uncertainty (e.g. 18) or a random age drawn from that interval (e.g. 19; 20). However, fossil age uncertainty can also be modelled explicitly by placing a prior on the fossil ages and co-estimating these along with other model parameters (21). Previous authors have demonstrated that different age interpretations can lead to substantial differences in empirical estimates of divergence times in analyses that directly incorporate fossils into the tree (22). In this paper we explore fossil age uncertainty as a potential source of error in FBD analyses using simulated and empirical data, and we describe how various methods of handling age uncertainty can affect the results. Our simulations show that fixing specimen ages can lead to erroneous estimates of divergence times but that incorporating stratigraphic age uncertainty explicitly using a hierarchical modelling approach substantially increases the chances of recovering the correct node ages. An analysis of Cetacea (the clade containing dolphins and whales) reveals that alternative approaches to handling specimen ages have major implications for dating speciation events within a historical context.

## 2 Methods

### 2.1 Simulated datasets

Our simulation study is based on the class Mammalia, which is very well represented in both molecular and paleontological databases. Mammals and their subgroups have also been the subject of a large number of molecular dating studies (23; 24; 25), as well as studies that rely on time-scaled phylogenies (26; 27). Simulated age uncertainty was based on the fossil record of North American mammals, which is relatively complete, has been studied in detail and is stratigraphically well constrained. Thus, the degree of age uncertainty incorporated into our simulations represents a best-case scenario for inferring the divergence times of mammal-like groups.

### 2.1.1 Simulation of extant species phylogenies and fossil samples

Trees were simulated under a birth-death process using the R package TreeSim (28). The speciation rate was set to λ = 0.15/Myr and the extinction rate to *µ* = 0.1/Myr, in accordance with estimates from (26) for the mammalian phylogeny. The birth-death process was stopped once *n_extant_* = 25 extant species had been reached. At this stage we rejected trees whose origin was not between 40 Ma and 100 Ma. This interval broadly encompasses the estimated origin time for many major groups of mammals (27; 24), but avoids simultaneously conditioning on tree age and tip number, which can be problematic (29).

Fossils were then sampled on the complete phylogeny using the R package FossilSim, following a Poisson process with a constant rate. The fossilization rate was set to ψ = 0.2/Myr, based on estimates of fossil recovery rates among Cenozoic mammals (30). Note that under this process, any number of fossils can be sampled for a given species. We did not filter the fossils further, as the current implementation of the FBD model in BEAST2 is designed for specimen-level data. We rejected phylogenies with less than 4 or more than 125 sampled fossils. A minimum of 4 was chosen to ensure convergence when analyzing the simulated data, while a maximum of 125 was chosen to avoid generating trees with a large number of extinct samples relative to the number of extant samples (*n* = 25), which is not typical in divergence time studies. The sampled tree was then obtained by assuming complete extant species sampling (ρ = 1) and discarding all unsampled lineages.

### 2.1.2 Simulation of sequences

Sequences were simulated for the extant species using seq-gen (31) via the R package phyclust (32). We simulated sequences of length 2000 nucleotides under an HKY+Γ model with 5 rate categories, following the substitution model used in (33). We used a log-normal uncorrelated clock, where the substitution rate for each branch of a given tree was sampled from a log-normal distribution with standard deviation 0.1. The mean of the log-normal distribution was drawn separately for each tree from a gamma distribution. A full list of all parameters used in the sequence simulation can be found in Table 1.

**Table 1:** List of all parameter values used to simulate sequences

| Parameter | Value |
| --- | --- |
| Mean substitution rate | $\sim \text{Gamma}(\alpha = 2, \beta = 2)/100$ |
| Branch-specific substitution rate | $\sim \text{LogNormal}(\text{mean rate}, 0.1)$ |
| Number of rate categories | 5 |
| Shape of the gamma distribution on rates ($\alpha$) | 0.25 |
| Transition/transversion ratio ($\kappa$) | 5 |
| Base frequencies | 0.25, 0.25, 0.25, 0.25 |

### 2.1.3 Simulation of fossil age uncertainty

The minimum and maximum age of each fossil occurrence was simulated based on the sampling interval ages of North American mammals. These intervals were downloaded from the Paleobiology Database (PBDB) on December 12th, 2017, using the following parameters: time intervals = Mesozoic and Cenozoic, region = North America, scientific name = Mammalia. If a fossil age could be assigned to multiple intervals, a single interval was selected at random by weighting all possible intervals by their frequency of appearance in the PBDB data. If no intervals appeared in the PBDB data for a simulated fossil, a random interval of fixed length was drawn. This length was fixed to the average length of all intervals present in the PBDB data, i.e. 8 Myr. Thus, the simulated interval for each fossil always included the correct age of the fossil. A visual representation of all the intervals used for simulation is shown in Figure 1.

**Figure 1:**
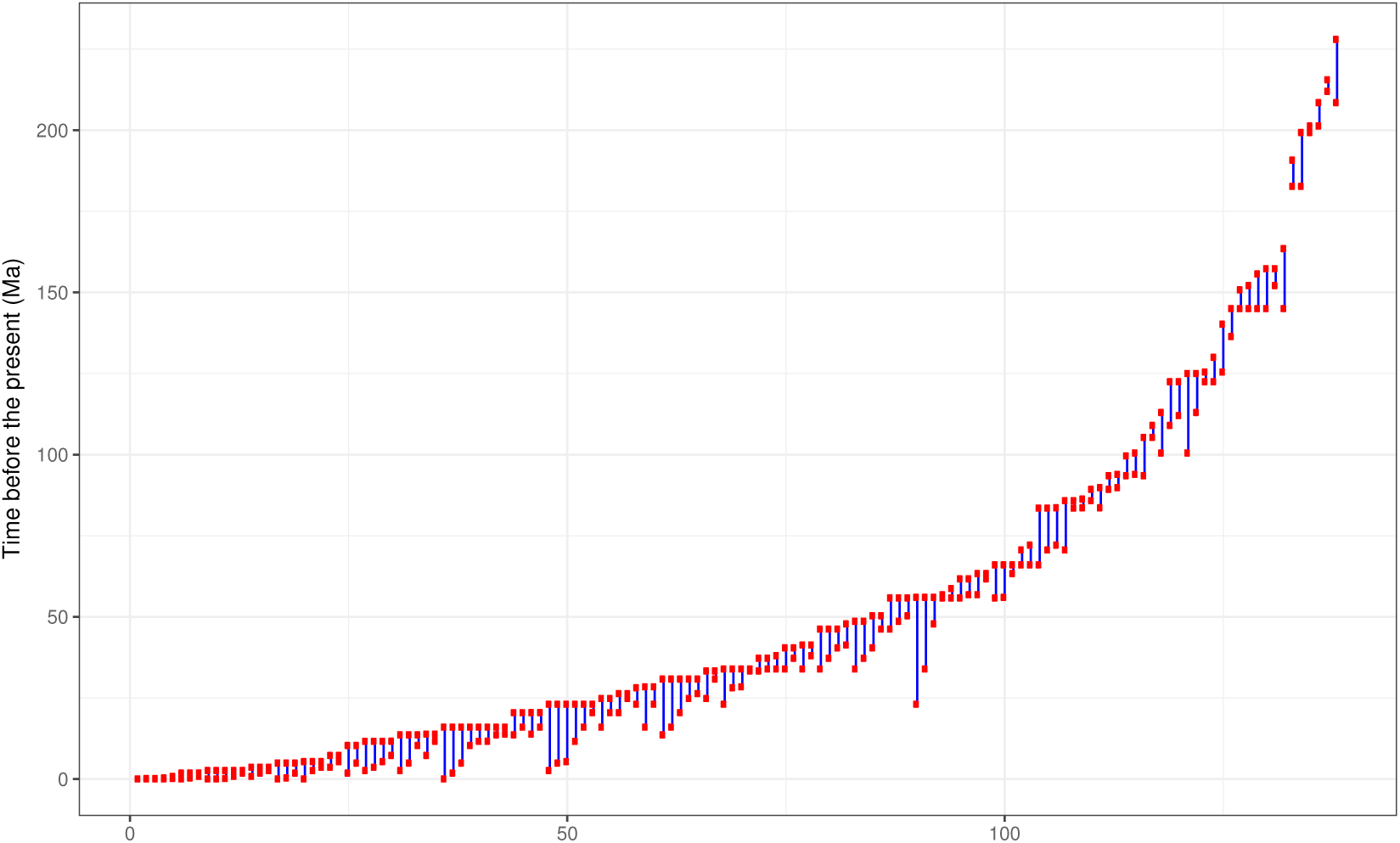
Representation of the age intervals obtained from PBDB for North American mammals sampled during the Mesozoic and Cenozoic. Intervals are ordered by the maximum age of the range, from youngest to oldest.

### 2.1.4 Bayesian inference

We used the Sampled Ancestors package (34), which provides an implementation of the fossilized birth-death process for the Markov Chain Monte-Carlo (MCMC) software BEAST2 (35), to perform Bayesian phylogenetic inferences on our simulated datasets.

The fossil ages were handled using five different methods, detailed here:

**Correct ages**: the fossil ages are fixed to the true ages as simulated.
**Interval ages**: the fossil ages are not fixed, but are sampled along with the other parameters within the simulated age range.
**Median ages**: the fossil ages are fixed to the midpoint of their simulated age range.
**Random ages**: the fossil ages are fixed to an age sampled uniformly at random inside of their simulated age range.
**Symmetric interval ages**: the fossil ages are not fixed, but are sampled along with the other parameters. Each fossil age is sampled within a symmetric interval around the true age of the fossil. The purpose of this setting was to evaluate the performance and accuracy in the situation where the midpoint of each prior interval was equal to the correct age.

A schematic representation of these different methods can be seen in Figure 2. Note that for the interval age methods, we sample trees as in (21). In particular, we set the probability density of the proposed tree to the FBD probability density if all fossil ages are within their intervals, and 0 otherwise. The effective prior on fossil ages is thus not a uniform prior, as the FBD model already induces a distribution on fossil ages.

**Figure 2:**
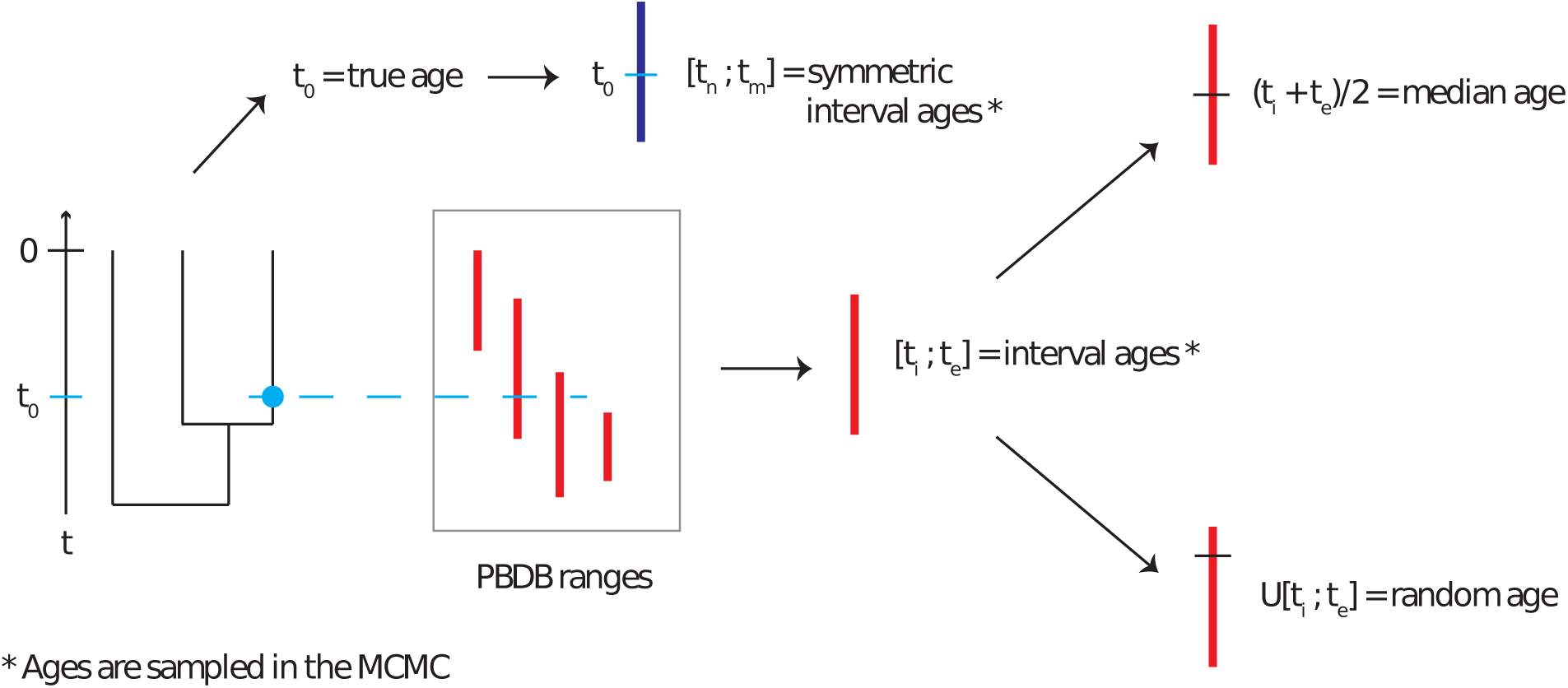
Representation of the age uncertainty simulation process. Phylogenies with fossils are simulated according to a birth-death-fossilization process. The correct age of each fossil is used to draw an age interval for that fossil from the set obtained from PBDB. This age interval is then used as the basis for the median and random age assignment. A symmetric age interval is also drawn from the correct age.

In all inferences, fossils were constrained to clades according to the correct tree topology. No sequence data was included for the fossil samples. The substitution model and clock model were set to the models used during simulation, and the priors used can be found in the Supplementary Materials.

### 2.2 Empirical dataset

To explore the impact of stratigraphic age uncertainty on empirical estimates of divergence times, we compiled a dataset of Cetacea containing both fossil occurrences and an alignment of sequences for extant species. This group was chosen based on the availability of a large molecular alignment representing almost all extant species, in combination with well-curated and comprehensive stratigraphic occurrence data. This group has also been the focus of a large number of molecular dating studies (36; 37; 38; 39; 40).

#### 2.2.1 Fossil occurrence data

We obtained data on 4473 fossil occurrences from the PBDB on April 5th, 2018, using the parameter “scientific name = Cetacea”. The full dataset could not be used due to mixing issues, so we subsampled 10% of the fossils at random, obtaining 448 fossil occurrences. The classification of taxa into suprageneric ranks was largely based on (41). A list of genera and their taxonomic membership as used in the subsample is provided in the Supplementary Materials. We used the minimum and maximum age for each fossil occurrence as recorded in the Paleobiology Database.

#### 2.2.2 Alignment

We used the alignment provided by (36), which contains sequences for 6 mitochondrial and 9 nuclear genes for 87 of 89 extant cetacean species. We excluded from our analysis the 3 outgroup taxa which were present in the original alignment, as our dataset contains no fossils for these families. The full alignment thus contains 87 sequences with 16,175 characters each.

Following (36), we divided this alignment into 28 partitions. The best substitution models for each partition were selected using bModelTest (42). A complete list of the substitution models used can be found in the Supplementary Materials.

#### 2.2.3 Bayesian inference

Bayesian phylogenetic inference was performed using the same model parameters and priors as for the simulated data, with the exception of the substitution model, which was set as specified in the previous section. As the correct ages of the fossils in this dataset are unknown, we limited our comparison to median ages, random ages and interval ages.

Topological constraints were set at both the genus and the family level, following the classification from PBDB, so that each genus or family formed a monophyletic clade in the tree. Samples whose position could not be determined were not included in any clade constraint, and thus could appear anywhere outside of the determined clades.

Following the model described in (43) for the analysis of empirical data, we unlinked the substitution models and among-site rate variation across partitions but linked the clock model and applied partition-specific rate multipliers to account for variation in evolutionary rates.

### 2.3 Data availability

All analyses, pre- and post-processing of the datasets were done using custom R scripts. These scripts and the XML configuration files used to run BEAST2 are included in the Supplementary Materials.

## 3 Results

### 3.1 Simulated datasets

#### 3.1.1 Accuracy

We measured the relative error of the median estimates for the divergence times and FBD model parameters obtained using different approaches to handling stratigraphic age uncertainty. We also calculated the coverage, i.e the number of analyses (out of 100 trees) in which the true parameter value was included in the 95% highest posterior density (HPD) interval. The error and coverage of the divergence times for each tree were averaged across all nodes of the extant tree, i.e. all nodes that were the most recent common ancestors (MRCA) of extant tips. Results for the divergence times, diversification rate and turnover estimates are shown in Figure 3.

**Figure 3:**
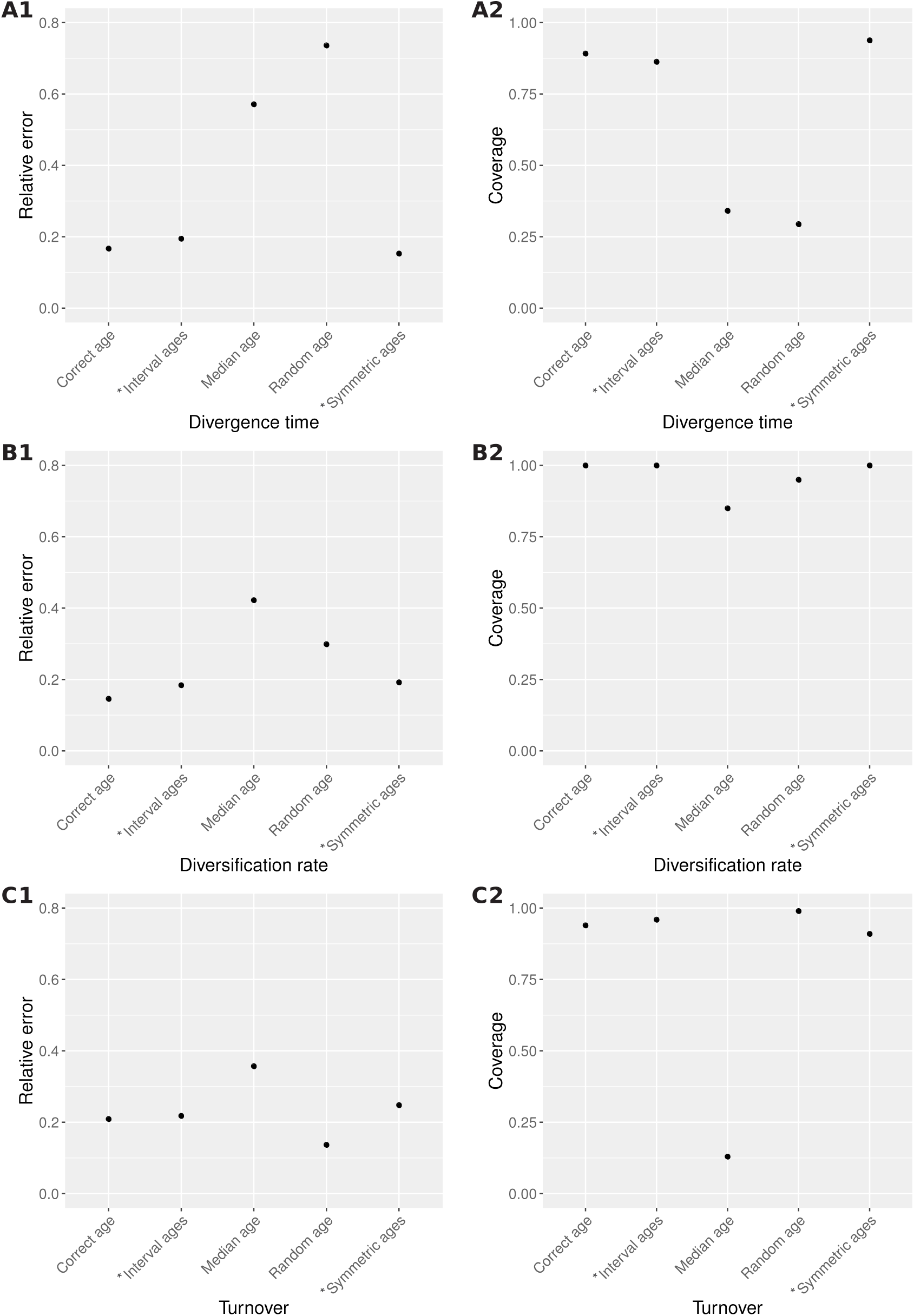
Average relative error of the median estimate (column 1) and 95% HPD coverage (column 2) achieved by different age handling methods for the following parameters: divergence times of extant species (A), diversification rate (B), and turnover (C). Ages sampled as part of the MCMC are marked by (*).

Using either interval ages or symmetric interval ages lead to error and coverage values that are very close to the results obtained using the correct age of the fossils. However, median and random ages did much worse than other methods. This is particularly apparent for estimates of divergence times, where error values range from 0.15 to 0.20 for correct, interval and symmetric interval ages, versus 0.57 and 0.74 for median and random ages, respectively. Similarly, the coverage of divergence times is 0.89 and 0.86 for correct and interval ages, respectively, versus 0.34 and 0.29 for median and random ages, respectively. The diversification rate is less sensitive to the choice of fossil ages, but still shows a relative error of 0.42 and a coverage of 0.85 for median ages, compared with 0.18 and 1.0 for interval ages. Median ages result in inaccurate estimates of turnover, with a coverage of 0.13, compared to values above 0.9 for all other methods. Overall, there are important discrepancies between the results obtained using correct ages, interval ages and symmetric interval ages, versus median ages and random ages.

The relative 95% HPD interval widths are shown in Table 2. Sampling fossil ages along with other model parameters (based on the PBDB or symmetric age intervals) did not result in wider HPD intervals than fixing the fossil ages to the truth. Fixing fossil ages to the wrong values (i.e using median ages or random ages) did not have a consistent effect on HPD interval width. For example, HPD intervals were wider for the divergence time estimates but narrower for the diversification rate. This reveals that the better coverage obtained for interval ages compared to median or random ages were not obtained at the expense of precision in the case of divergence times, i.e higher coverage is not simply due to wider HPD intervals.

**Table 2:**
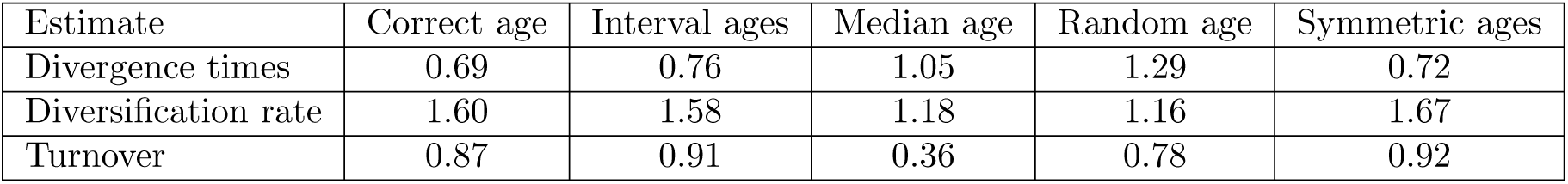
Average relative width of the 95% HPD intervals obtained for different estimates using different methods. The relative width is calculated as the width of the HPD interval divided by the true value.

#### 3.1.2 Performance

We evaluated the performance of the different inference methods by calculating the processing time required per effective sample. We used the effective sample sizes for the posterior distribution and for the total height of the tree. The results are shown in Table 3. We observed no clear correlation between sampling or fixing fossil ages and the performance in this dataset.

**Table 3:** Average performance of the MCMC using different age handling methods. The performance is calculated as the CPU time (in seconds) per effective sample of the posterior distribution and of the total length of the tree.

| time/ESS | Correct age | Median age | Interval ages | Random age | Symmetric ages |
| --- | --- | --- | --- | --- | --- |
| Posterior | 188.2 | 36.8 | 163.9 | 131.7 | 150.4 |
| Total length | 12.7 | 15.8 | 22.1 | 26.0 | 26.2 |

### 3.2 Empirical dataset

Figure 4 shows the MCC trees obtained using different methods of fixing ages, restricted to extant tips. There are few differences in topology, which is expected as we applied strong topological constraints in this analysis. However, divergence time estimates vary considerably between different approaches to handling age uncertainty, and is most apparent for older nodes in the tree. For instance, the most recent common ancestor (MRCA) of all extant cetaceans is 44 Ma old using interval ages, in contrast to 50 and 61 Ma using random and median ages, respectively. The relative difference between the median node ages inferred with interval ages versus median ages, averaged across all nodes, is 15%. However, there is wide variability; some nodes show almost no difference (< 1%), while in other cases the median age estimate obtained using median ages is double the estimate obtained using interval ages. The relative length of the 95% HPD intervals for the divergence time estimates is 52% of the median estimate for interval ages, 54% for median ages and 26% for random ages, also averaged across all nodes. Thus using random ages lead to much narrower posterior distributions for the divergence time estimates.

**Figure 4:**
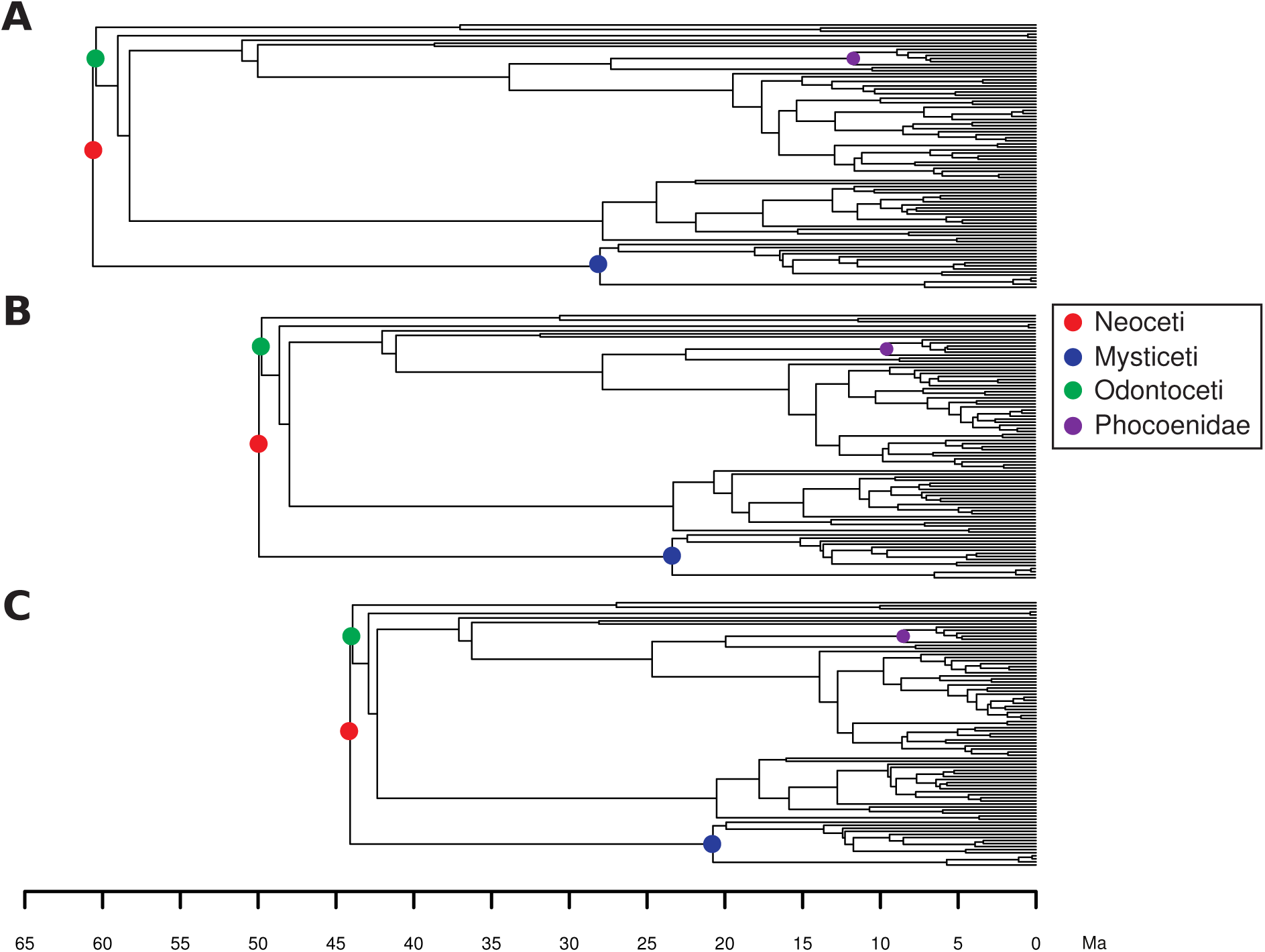
MCC trees inferred for the Cetacea dataset using the FBD process with fossil ages fixed to median ages (A), random ages (B) or sampled within the known interval of uncertainty (C). The major clades and the clade shown in Figure 5 are highlighted.

An example of the strong influence exerted by fixing fossil ages on estimated node ages is shown in Figure 5. We can see that the posterior distribution obtained using interval ages is much wider. However, when using median or random ages, the age of the node is constrained to within a much narrower interval. The fossil specimen imposes a lower bound on the distribution that is potentially in conflict with the phylogenetic data and/or other age constraints, resulting in a posterior distribution with a strong peak at the lower bound. For this node, the 95% HPD interval is of length 2.92 for the interval ages, 3.91 for median ages and 2.65 for random ages.

**Figure 5:**
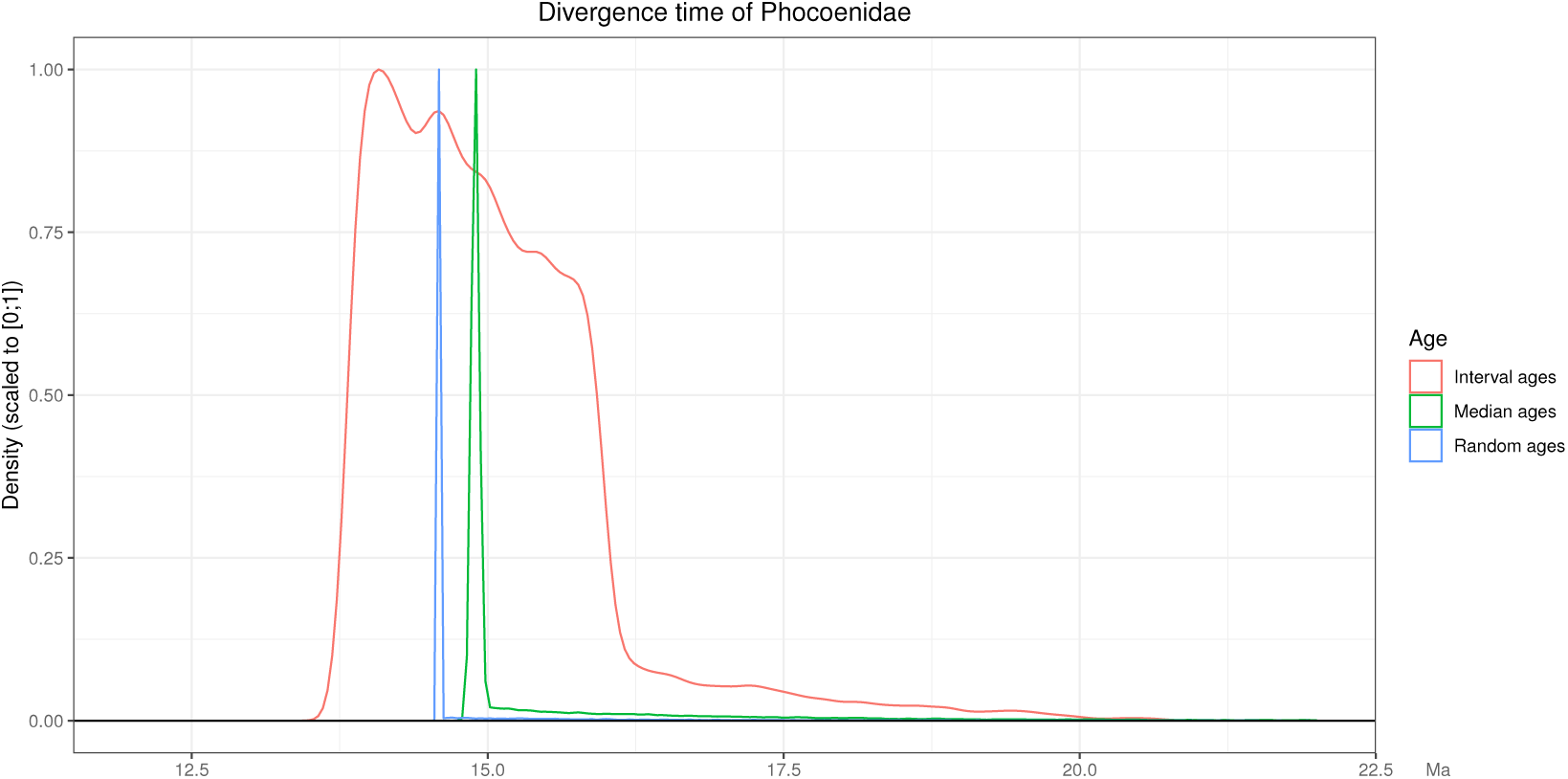
Posterior density obtained for the most recent common ancestor of the family Phocoenidae in the Cetacea dataset using the FBD process with fossil ages fixed to median ages, random ages or sampled within the known interval of uncertainty. The densities were scaled to the interval [0; 1].

Figure 6 shows a comparison of the estimates of the FBD process parameters obtained using different methods of age handling. For these parameters, all estimates show a trend going from median to random to interval ages. The diversification rate is most robust to the choice of fossil age, as all the HPD intervals overlap. However, we see a trend for an increasing diversification rate estimate, from median to random to interval ages. The turnover is estimated to be higher and the sampling proportion much lower with median ages than with interval ages. These trends in parameter estimates are likely to be in part correlated to the observed differences in divergence time estimates. For example, as the number of fossil samples increases with the age of the process and with the sampling rate, estimating a greater height for the tree leads to lower estimates of sampling proportion and vice-versa. Similarly, as the number of extant species increases with the age of the process and with the diversification rate, estimating a greater height for the tree leads to lower estimates of diversification rates and vice-versa.

**Figure 6:**
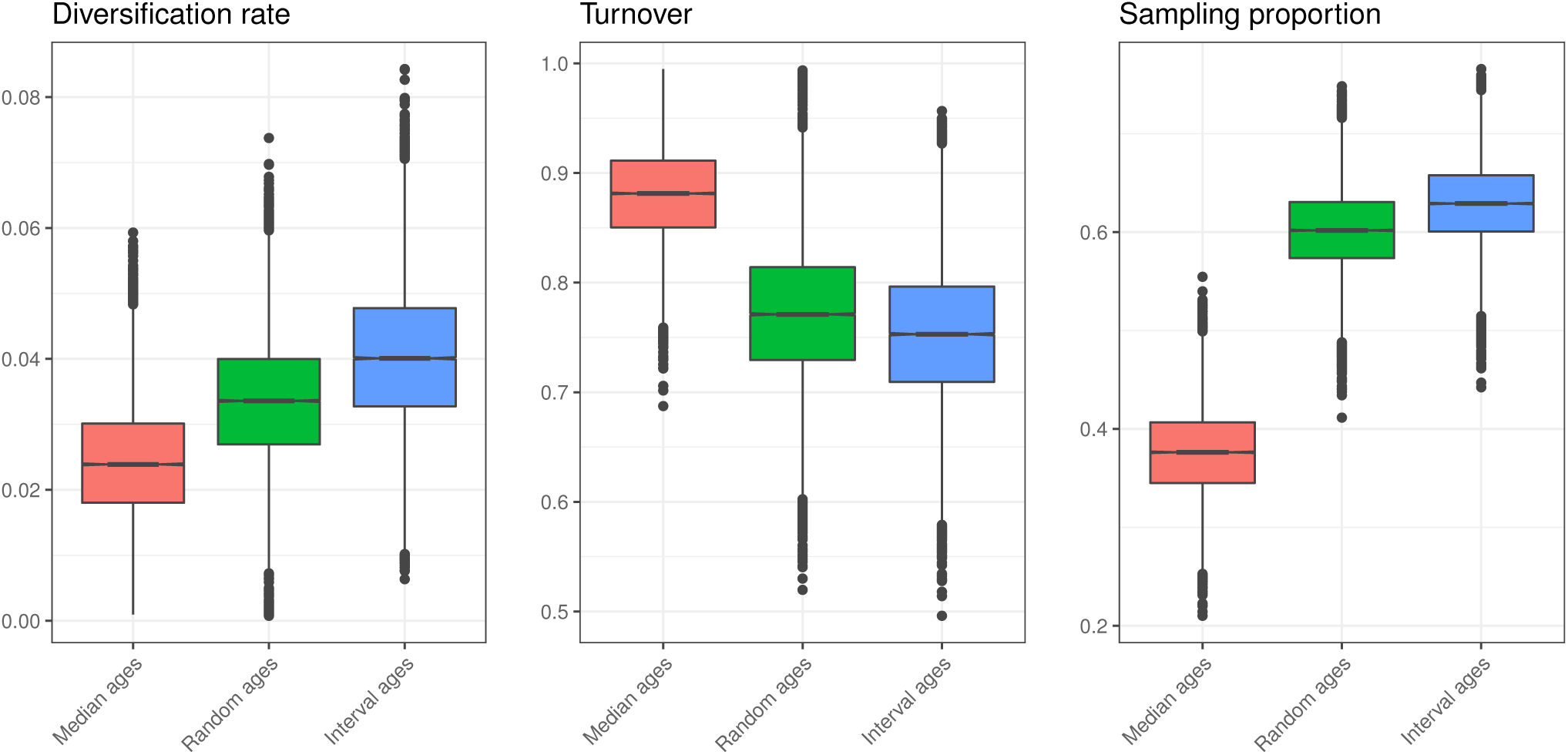
Estimates of the diversification rate, turnover and sampling proportion obtained for the Cetacea dataset using the FBD process with fossil ages fixed to median ages, random ages or sampled within the known interval of uncertainty.

## 4 Discussion

The age of fossils is not known precisely and instead is associated with a range of uncertainty that results from the nature of the geological record. Our simulations demonstrate that the popular approach of fixing fossil ages to point estimates within the known interval of uncertainty in molecular dating analysis can led to erroneous estimates of node ages. Using this strategy <35% of node ages were recovered within the Bayesian 95% HPD intervals. Conversely, explicitly incorporating fossil age uncertainty led to substantially more accurate estimates of node ages and other model parameters. It may be tempting to fix the age of fossils to reduce the computational burden. However, our results based on both the simulated and empirical datasets show that co-estimating fossil ages does not necessarily incur additional computational cost. Unless researchers have extremely good motivation for doing so, fossil ages should not be fixed in divergence time analysis that directly incorporate fossils into the tree. Emphasis on establishing rigorous and transparent calibration information that evolved in the context of node dating should also apply to other approaches to calibration, including inference under the FBD process (17; 44; 22).

The analysis of the cetacean dataset illustrates how strongly estimates of both divergence times and biologically relevant parameters, such as diversification rate, can be affected by the choices made in handling fossil age uncertainty. The difference between the median posterior estimate obtained for the age of Neoceti (crown cetaceans) using fixed median ages versus interval ages is nearly 20 Ma. Furthermore, analysing this dataset with median ages would show a strong discrepancy between the divergence time estimates obtained using the FBD model and the estimates obtained using fossil calibration in the original node dating analysis (36), which estimated the origin of the Neoceti to be 36 Ma. However, accounting for stratigraphic age uncertainty shows that this is not the case: the MRCA age obtained using interval ages matches more closely both with the original analysis and with more recent studies such as (39), which estimated the MRCA age to be 39 Ma. It is not possible to definitively determine which outcome is closer to the truth in our empirical analysis, but our simulations clearly indicate that estimates obtained using interval ages should be considered the most reliable.

It is worth noting that the average range of age uncertainty associated with fossils included in our simulations and empirical analysis is relatively small (8 and 4 Myr, respectively). This reflects our decision to focus on well-studied Cenozoic fauna with extant representatives, but the fossil record of many taxonomic groups and time periods will be associated with much greater uncertainty. For example, the age of many pre-Cenozoic deposits are poorly constrained. Thus, the discrepancies obtained using different age handling methods have the potential to be much larger than those demonstrated in this study. This may be especially important to consider in the context of FBD analyses for groups that have no extant representatives.

In our experiments, no character data was included for extinct samples, meaning fossil recovery times can inform the FBD model parameters, but the phylogenetic position of these samples cannot be estimated. If morphological character data is available for fossils, their phylogenetic position can also be inferred along with divergence times (43; 45), meaning no taxonomic decisions have to be made *a priori* by assigning fossils to clades as done here. This could increase the impact of mishandling fossil age uncertainty, especially since morphological character matrices available for fossils are typically small (e.g. 50-200 characters).

Finally, it is worth emphasising the distinction between the age range associated with fossil specimens and fossil species. The latter is known as the stratigraphic range of a species, and represents the interval between the first and last appearance times. Here, we implemented the specimen-based FBD process, meaning all available specimens were incorporated into the analysis. Although we note some studies have applied this model to the analysis of stratigraphic range data, this is not technically appropriate. Instead, stratigraphic ranges should be analysed under the FBD range process (46), however no implementation in BEAST2 is yet available. We note that when this model does become available, the uncertainty associated with specimens representing the ends of stratigraphic ranges should be incorporated into the analysis, rather than being fixed, otherwise we anticipate similar performance issues to those demonstrated in this study.

## 5 Conclusions

In this study we demonstrate that the choice of method for handling fossil age uncertainty can have important effects on estimates of species divergence times obtained under the FBD process. Our simulation results clearly favour a Bayesian hierarchical approach to handling fossil age uncertainty based on the actual age intervals, as opposed to fixing the ages to an arbitrary value inside that interval. In addition, our empirical dataset demonstrates that the rigid age constraints given by fixed fossil ages can lead to age estimates that are very different from those obtained using a traditional node dating approach, whereas a more flexible approach to handling fossil ages recovers similar estimates. Thus we strongly recommend against fixing fossil ages in FBD analyses. Overall, this work illustrates the critical importance of accurately reflecting the available information regarding uncertainty in Bayesian phylogenetic analyses. As we demonstrate, discarding this information can have detrimental impacts on the accuracy of the results.

## Supporting information

## 6 Acknowledgements

We thank Chi Zhang for advice in setting up the analysis of empirical data. We also thank Mark D. Uhen for keeping the cetacean data in the PBDB up to date. JBS is supported in part by the European Research Council under the Seventh Framework Programme of the European Commission (PhyPD: grant agreement number 335529). RCMW is funded by the ETH Zörich Postdoctoral Fellowship and Marie Curie Actions for People COFUND programme.

