## Supplementary material for "Ignoring stratigraphic age uncertainty leads to erroneous estimates of species divergence times under the fossilized birth-death process": taxonomy.pdf

**Electronic Supplementary Material Table 1. Taxonomic membership of the genera in the empirical dataset of Cetacea.**

| Genus | Family | Reference |
|---|---|---|
| *Acrophyseter* | Physeteroidea | [1] |
| *Africanacetus* | Ziphiidae | [1] |
| *Aglaocetus* | Chaeomysticeti | [1] |
| *Agorophius* | Agorophiidae | [1] |
| *Amphicetus* | Cetacea | |
| *Ancalecetus* | Basilosauridae | [1] |
| *Andrewsiphius* | Remingtonocetidae | [1] |
| *Aporotus* | Ziphiidae | [1] |
| *Argyrocetus* | Eurhinodelphinidae | [1] |
| *Astadelphis* | Delphinidae | [1] |
| *Atocetus* | Pithanodelphininae | [1] |
| *Attockicetus* | Remingtonocetidae | [1] |
| *Babiacetus* | Protocetidae | [1] |
| *Balaena* | Balaenidae | [1] |
| *Balaenoptera* | Balaenopteridae | [1] |
| *Balaenotus* | Cetacea | |
| *Balaenula* | Balaenidae | [1] |
| *Basilosaurus* | Basilosauridae | [1] |
| *Basilotritus* | Basilosauridae | [1] |
| *Belemnoziphius* | Ziphiidae | [2] |
| *Berardius* | Ziphiidae | [1] |
| *Brandtocetus* | Cetotheriidae | [3] |
| *Caperea* | Neobalaenidae | [1] |
| *Cetotheriophanes* | Balaenopteridae | [4] |
| *Cetotherium* | Cetotheriidae | [1] |
| *Chilcacetus* | Eurhinodelphinidae | [1] |
| *Coronodon* | Mysticeti | [5] |
| *Dalanistes* | Remingtonocetidae | [1] |
| *Delphinapterus* | Monodontidae | [1] |
| *Delphinodon* | Kentriodontinae | [1] |
| *Delphinus* | Delphinidae | [1] |
| *Denebola* | Monodontidae | [1] |
| *Dorudon* | Basilosauridae | [1] |
| *Eschrichtius* | Eschrichtiidae | [1] |
| *Eubalaena* | Balaenidae | [1] |
| *Eucetotherium* | Cetotheriidae | [6] |
| *Eurhinodelphis* | Eurhinodelphinidae | [1] |
| *Feresa* | Delphinidae | [1] |
| *Globicephala* | Delphinidae | [1] |
| *Globicetus* | Ziphiidae | [1] |
| *Goniodelphis* | Iniidae | [1] |
| *Grampus* | Delphinidae | [1] |

| Genus | Family | Reference |
|---|---|---|
| *Hadrodelphis* | Lophocetinae | [1] |
| *Hemisyntrachelus* | Delphinidae | [1] |
| *Herpetocetus* | Cetotheriidae | [1] |
| *Heterocetus* | Cetacea | |
| *Hoplocetus* | Physeteroidea | [1] |
| *Hyperoodon* | Ziphiidae | [1] |
| *Ichthyolestes* | Pakicetidae | [1] |
| *Indopacetus* | Ziphiidae | [1] |
| *Kentriodon* | Kentriodontinae | [1] |
| *Kharodacetus* | Protocetidae | [1] |
| *Kogia* | Kogiidae | [1] |
| *Lagenodelphis* | Delphinidae | [1] |
| *Lagenorhynchus* | Delphinidae | [1] |
| *Masracetus* | Basilosauridae | [1] |
| *Mauicetus* | Chaeomysticeti | [1] |
| *Megaptera* | Balaenopteridae | [1] |
| *Mesoplodon* | Ziphiidae | [1] |
| *Mithridatocetus* | Cetotheriidae | [6] |
| *Monodon* | Monodontidae | [1] |
| *Neophocaena* | Phocoenidae | [1] |
| *Neosqualodon* | Squalodontidae | [1] |
| *Ninjadelphis* | Allodelphinidae | [7] |
| *Ninoziphius* | Ziphiidae | [1] |
| *Notiocetus* | Cetacea | |
| *Ocucajea* | Basilosauridae | [1] |
| *Orcaella* | Delphinidae | [1] |
| *Orcinus* | Delphinidae | [1] |
| *Pachyacanthus* | Delphinida | [8] |
| *Palaeophocaena* | Cetacea | |
| *Parapontoporia* | Lipotidae | [1] |
| *Parietobalaena* | Chaeomysticeti | [1] |
| *Patriocetus* | Patriocetidae | [1] |
| *Peponocephala* | Delphinidae | [1] |
| *Phocoena* | Phocoenidae | [1] |
| *Physeterula* | Physeteridae | [1] |
| *Pinocetus* | Balaenopteroidea | [1] |
| *Piscolithax* | Phocoenidae | [1] |
| *Platyosphys* | Cetacea | |
| *Plesiocetopsis* | Tranatocetidae | [3] |
| *Plesiocetus* | Cetacea | |
| *Pomatodelphis* | Platanistidae | [1] |
| *Pseudorca* | Delphinidae | [1] |
| *Sachalinocetus* | Waipatiidae | [9] |
| *Saurocetes* | Iniidae | [1] |

| Genus | Family | Reference |
|---|---|---|
| *Scaldicetus* | Physeteroidea | [1] |
| *Schizodelphis* | Eurhinodelphinidae | [1] |
| *Simocetus* | Simocetidae | [1] |
| *Sitsqwayk* | Mysticeti | [10] |
| *Sousa* | Delphinidae | [1] |
| *Squalodon* | Squalodontidae | [1] |
| *Stenella* | Delphinidae | [1] |
| *Stromerius* | Basilosauridae | [1] |
| *Thinocetus* | Chaeomysticeti | [1] |
| *Tohoraata* | Eomysticetidae | [1] |
| *Tokarahia* | Eomysticetidae | [1] |
| *Tursiops* | Delphinidae | [1] |
| *Tusciziphius* | Ziphiidae | [1] |
| *Xiphiacetus* | Eurhinodelphinidae | [1] |
| *Zarhachis* | Platanistidae | [1] |
| *Ziphiodelphis* | Eurhinodelphinidae | [1] |
| *Ziphirostrum* | Ziphiidae | [1] |
| *Ziphius* | Ziphiidae | [1] |
| *Zygophyseter* | Physeteroidea | [1] |
| *Zygorhiza* | Basilosauridae | [1] |
