## Supplementary material for "Ignoring stratigraphic age uncertainty leads to erroneous estimates of species divergence times under the fossilized birth-death process": priors_cetaceans_beast2.pdf

### List of substitution models and priors used for the cetaceans dataset.

| Partition | Substitution model | Invariant proportion |
|---|---|---|
| Alb 12 | HKY + $\Gamma$ + I | 0.32 |
| Alb 3 | TN93 + $\Gamma$ + I | 0.22 |
| Bdnf 12 | TN93 + $\Gamma$ + I | 0.96 |
| Bdnf 3 | TN93 + $\Gamma$ + I | 0.24 |
| Cmos 12 | GTR + $\Gamma$ + I | 0.31 |
| Cmos 3 | HKY + $\Gamma$ + I | 0.20 |
| Co1 12 | GTR + $\Gamma$ + I | 0.76 |
| Co1 3 | TN93 + $\Gamma$ | |
| Cytb 12 | HKY + $\Gamma$ + I | 0.54 |
| Cytb 3 | GTR + $\Gamma$ + I | 0.16 |
| Irbp 12 | HKY + $\Gamma$ + I | 0.53 |
| Irbp 3 | GTR + $\Gamma$ + I | 0.19 |
| Nd3 12 | TN93 + $\Gamma$ + I | 0.36 |
| Nd3 3 | TN93 | |
| Nd4 12 | TN93 + $\Gamma$ + I | 0.38 |
| Nd4 3 | HKY + $\Gamma$ | |
| Nd4l 12 | TN93 + $\Gamma$ + I | 0.31 |
| Nd4l 3 | K81 | |
| Noncoding | GTR + $\Gamma$ | |
| Prion 12 | GTR + $\Gamma$ + I | 0.23 |
| Prion 3 | GTR + $\Gamma$ + I | 0.17 |
| Rag2 12 | TN93 + $\Gamma$ + I | 0.17 |
| Rag2 3 | HKY + $\Gamma$ + I | 0.18 |
| Rrna | GTR + $\Gamma$ + I | 0.35 |
| Sry 12 | GTR | |
| Sry 3 | GTR | |
| Tbx4 12 | TVM + $\Gamma$ + I | 0.42 |
| Tbx4 3 | K81 + $\Gamma$ + I | 0.28 |

| Parameter | Prior distribution |
|---|---|
| FBD diversification rate | Uniform$[0; \infty[$ |
| FBD turnover | Uniform$[0; 1[$ |
| FBD sampling proportion | Uniform$[0; 1[$ |
| Origin of the FBD process | Uniform$[0; \infty[$ |
| HKY $\kappa$ parameter | LogNormal(1.0,1.25) |
| TN93 $\kappa_1$, $\kappa_2$ parameter | LogNormal(1.0,1.25) |
| GTR AC, AT, CG, GT rates | Gamma($\alpha = 0.05$,$\beta = 10.0$) |
| GTR AG rate | Gamma($\alpha = 0.05$,$\beta = 20.0$) |
| Shape of the $\Gamma$ distribution | Exponential(1.0) |
| Linked clock rate | Uniform$[0; \infty[$ |
