## Supplementary material for "Ignoring stratigraphic age uncertainty leads to erroneous estimates of species divergence times under the fossilized birth-death process": README.txt

This folder contains the data related to the empirical cetaceans dataset.

The alignments subfolder contains the original sequence alignments obtained from Steeman et al. 2009.
The XML subfolder contains the xml configuration files used to run FBD analyses on this dataset with interval ages, random ages or median ages.

The fossils subfolder contains the data used to integrate fossil occurrence data in the analysis:
- the file cetacea_pbdb_ss0.1.csv shows the PBDB record for all fossils used in the analysis.
- the file taxonomy.pdf contains the taxonomic classification of genera into families. 
- the file families.tre shows the topology of families (obtained from Marx et al. 2016) used to create the topological constraints.

The file priors_cetaceans_beast2.pdf contains the list of substitution models and priors used to run BEAST2 on this dataset.
