## Supplementary material for "Ignoring stratigraphic age uncertainty leads to erroneous estimates of species divergence times under the fossilized birth-death process": mammals_accuracy.R

```
# analyze error and coverage of parameters from BEAST2 logfiles
# only divergence times on which a taxonomic constraint was present appear here
mammals_log_analysis = function(treefile, outfolder) {
  library(ape)
  library(coda)
  library(Metrics)
  
  spec_rate = 0.15
  ext_rate = 0.1
  sampl_rate = 0.2
  nextant = 25
  
  load(treefile)
  
  names = c("correct_ages", "median_ages", "interval_ages", "random_ages", "norm_intvl_ages")
  FBDp_names = c("diversificationRateFBD","turnoverFBD", "samplingProportionFBD", "SACountFBD")
  FBDp_names = c(FBDp_names, "rate.mean")
  
  trueFBDp = c(spec_rate - ext_rate, ext_rate / spec_rate, sampl_rate / (ext_rate + sampl_rate), 0, 0)
  names(trueFBDp) = FBDp_names
  
  analysis = list()
  ESS = list(npb = 0, lpb = c(), whys = list())
  
  for(i in 1:length(trees)) {
    print(i)
    logs = list()
    for(x in 1:length(names)) {
      logs[[x]] = read.table(file.path(outfolder, paste0("FBD_", names[[x]],"_",i,".log")),header = T,comment.char = "#")
      n = nrow(logs[[x]])
      logs[[x]] = logs[[x]][(round(n/10)+1):n,]
    }
    
    why = c()
    tree = samp_trees[[i]]
    clades = .find_clades(tree, nextant)
    descend = .get_desc(tree)
    ages = n.ages(tree)
    
    #true substitution rate & SA count
    trueFBDp['rate.mean'] = rates[i]
    trueFBDp['SACountFBD'] = 0
    for(tip in 1:length(tree$tip.label)) {
      if(tree$edge.length[which(tree$edge[,2] == tip)] == 0) trueFBDp['SACountFBD'] = trueFBDp['SACountFBD'] +1
    }
    
    for(x in 1:length(names)) {
      for(y in FBDp_names) {
        #error on FBD parameters
        if(y == 'SACountFBD') {
          analysis[[paste0(names[x],".",y,".relative_error")]] = c(analysis[[paste0(names[x],".",y,".rel.error")]], abs(median(logs[[x]][,y]) - trueFBDp[y])/length(tree$tip.label))
        }
        else {
          analysis[[paste0(names[x],".",y,".relative_error")]] = c(analysis[[paste0(names[x],".",y,".abs.error")]], abs(median(logs[[x]][,y]) - trueFBDp[y])/trueFBDp[y])
          analysis[[paste0(names[x],".",y,".median")]] = c(analysis[[paste0(names[x],".",y,".median")]], median(logs[[x]][,y]))
        }
        HPD = coda::HPDinterval(as.mcmc(logs[[x]][,y]))
        analysis[[paste0(names[x],".",y,".coverage")]] = c(analysis[[paste0(names[x],".",y,".coverage")]], (trueFBDp[y] >= HPD[1]) && (trueFBDp[y] <= HPD[2]))
        analysis[[paste0(names[x],".",y,".HPD_width")]] = c(analysis[[paste0(names[x],".",y,".HPD_width")]], (HPD[2] - HPD[1])/trueFBDp[y] )
        
        #ESS checks on FBD pars
        ess = ESScalc(logs[[x]][,y])
        if(is.na(ess) || ess<200) why = c(why,paste0(names[x],".",y))
      }
      
      rel.error = c()
      cover = c()
      #for each clade previously identified -> error & coverage of div time
      for(c in clades) {
        true_t = ages[which(sapply(descend, identical, c))]
        y = paste0("mrcatime.", paste0(c,collapse = "."), ".")
        rel.error = c(rel.error, abs(true_t - median(logs[[x]][,y]))/true_t)
        HPD = HPDinterval(as.mcmc(logs[[x]][,y]))
        cover = c(cover, (true_t >= HPD[1] - 1e-6) && (true_t <= HPD[2] + 1e-6))
        
        #ESS checks on div times
        ess = ESScalc(logs[[x]][,y])
        if(is.na(ess) || ess<200) why = c(why,paste0(names[x],".",y))
      }
      #for one tree summarize coverage as %, error as mean relative error
      analysis[[paste0(names[x],".divtime.relative_error")]] = c(analysis[[paste0(names[x],".divtime.relative_error")]], mean(rel.error))
      analysis[[paste0(names[x],".divtime.coverage")]] = c(analysis[[paste0(names[x],".divtime.coverage")]], sum(cover)/length(cover))
    }
    
    #ESS check summary
    if(length(why) > 0) {
      ESS$npb = ESS$npb + 1
      ESS$lpb = c(ESS$lpb,i)
      ESS$whys[[ESS$npb]] = why
    }
  }
  
  for(x in 1:length(names)) {
    for(y in FBDp_names) {
      analysis[[paste0(names[x],".",y,".coverage")]] = sum(analysis[[paste0(names[x],".",y,".coverage")]])/length(trees)
    }
    for(y in c("diversificationRateFBD","turnoverFBD", "samplingProportionFBD")) {
      analysis[[paste0(names[x],".",y,".rmse")]] = Metrics::rmse(analysis[[paste0(names[x],".",y,".median")]], rep(trueFBDp[y], length(trees)))
    }
    analysis[[paste0(names[x],".rate.mean.rmse")]] = Metrics::rmse(analysis[[paste0(names[x],".rate.mean.median")]], rates)
  }
  
  analysis = lapply(analysis, function(x) {names(x) = NULL; x})
  analysis$ESS = ESS
  analysis
}

# analyze divergence times error and coverage based on trees file - all MRCAs of extant tips are considered
mammals_allnodes_divtimes = function(treefile, outfolder, tmp_file = "tmp_analysis.RData") {
  library(ape)
  library(coda)
  load(treefile)
  nextant = 25
  
  names = c("correct_ages", "median_ages", "interval_ages", "random_ages", "norm_intvl_ages")
  if(file.exists(tmp_file)) {
    load(tmp_file)
    analysis = get("analysis")
    initi = length(analysis[[1]])+1
  } else {
    analysis = list()
    initi = 1
  }
  if(initi > length(trees)) return(analysis)
  
  for(i in initi:length(trees)) {
    print(i)
    mrca_list = list(ages = c(), tips = list())
    used_nodes = c()
    ages = n.ages(samp_trees[[i]])
    used_tips = nextant
    
    all_mrca = mrca(samp_trees[[i]])
    for(t1 in 1:(used_tips-1)) {
      idx1 = which(rownames(all_mrca) == samp_trees[[i]]$tip.label[t1])
      for(t2 in (t1+1):used_tips) {
        idx2 = which(colnames(all_mrca) == samp_trees[[i]]$tip.label[t2])
        mrca = all_mrca[idx1,idx2]
        if(!mrca %in% used_nodes) {
          used_nodes = c(used_nodes, mrca)
          mrca_list$tips[[length(mrca_list$tips)+1]] = c(samp_trees[[i]]$tip.label[t1], samp_trees[[i]]$tip.label[t2])
          mrca_list$ages = c(mrca_list$ages, ages[mrca])
        }
      }
    }
    
    for (nm in names) {
      beast_trees = read.nexus(file.path(outfolder, paste0("FBD_", nm,"_",i,".trees")))
      # seriously, wtf, ape ?
      beast_trees = lapply(beast_trees, function(x) {x$tip.label = attr(beast_trees,"TipLabel"); x})
      n = length(beast_trees)
      beast_trees = beast_trees[(round(n/10)+1):n]
      
      offs = rep(0, length(beast_trees))
      
      tmp = list()
      for(mr in mrca_list$tips) tmp[[paste0(mr[1], ".", mr[2])]] = rep(0, length(beast_trees))
      for (bti in 1:length(beast_trees)) {
        ages = n.ages(beast_trees[[bti]])
        for(mr in mrca_list$tips) {
          t = ages[getMRCA(beast_trees[[bti]], mr)] + offs[bti]
          tmp[[paste0(mr[1], ".", mr[2])]][bti] = t
        }
      }
      rel_error = c()
      cover = wdth = c()
      for(j in 1:length(mrca_list$ages)) {
        rel_error = c(rel_error, abs(median(tmp[[paste0(mrca_list$tips[[j]][1], ".", mrca_list$tips[[j]][2])]]) 
                                     - mrca_list$ages[j])/mrca_list$ages[j])
        HPD = HPDinterval(as.mcmc(tmp[[paste0(mrca_list$tips[[j]][1], ".", mrca_list$tips[[j]][2])]]))
        cover = c(cover, (mrca_list$ages[j] >= HPD[1] - 1e-6) && (mrca_list$ages[j] <= HPD[2] + 1e-6))
        wdth = c(wdth, (HPD[2] - HPD[1])/mrca_list$ages[j])
      }
      
      analysis[[paste0(nm,".extmrca.divtime.relative_error")]] = c(analysis[[paste0(nm,".extmrca.divtime.relative_error")]], mean(rel_error))
      analysis[[paste0(nm,".extmrca.divtime.coverage")]] = c(analysis[[paste0(nm,".extmrca.divtime.coverage")]], sum(cover)/length(cover))
      analysis[[paste0(nm,".extmrca.divtime.HPD_width")]] = c(analysis[[paste0(nm,".extmrca.divtime.HPD_width")]], mean(wdth))
    }
    save(analysis,file=tmp_file)
  }
  #if(file.exists(tmp_file)) file.remove(tmp_file)
  analysis
}

# summarizes results across all trees dataset
mammals_summary = function(analysis) {
  res = list()
  if(!is.null(analysis$ESS)) {
    res$ESS = list(npb = analysis$ESS$npb, lpb = analysis$ESS$lpb)
    analysis$ESS = NULL
  }
  
  for(i in 1:length(analysis)) {
    if(length(analysis[[i]]) > 1) res[[names(analysis)[i]]] = mean(analysis[[i]])
    else res[[names(analysis)[i]]] = analysis[[i]]
  }
  res
}

ESScalc = function(values,burnin=0.1) {
  ntot = length(values)
  iS = floor(ntot*burnin)
  values = values[(iS+1):ntot]
  nS = length(values)
  av = mean(values)
  l = 0
  stp = F
  while(!stp && l<nS) {
    ssm = sum((values[1:(nS-l)]-av)*(values[(l+1):nS]-av))
    ssm = ssm/(nS-l)
    
    if(l==0) {
      AC = ssm
      ssm0 = ssm
    }
    else if(l%%2==1) ssm_tmp = ssm
    else {
      if(ssm+ssm_tmp >0) AC = AC + 2*(ssm_tmp+ssm)
      else stp = T
    }
    l=l+1
  }
  
  if(l==nS) {
    warning("ESS calculation reached max lag")
    ESS = Inf
  }
  else ESS = nS*ssm0/AC
  ESS
}

.get_desc = function(tree) {
  aux = function(node, result) {
    desc = which(tree$edge[,1] == node)
    if(length(desc) == 0) {
      result[[node]] = node
      return(result)
    }
    result = aux(tree$edge[desc[1],2], result)
    result = aux(tree$edge[desc[2],2], result)
    result[[node]] = c(result[[tree$edge[desc[1],2]]], result[[tree$edge[desc[2],2]]])
    
    return(result)
  }
  
  aux(length(tree$tip.label) + 1, list())
}

n.ages = function(tree){
  
  node.ages <- TreeSim::getx(tree, sersampling = 1)[1:(tree$Nnode+length(tree$tip))]
  names(node.ages) <- 1:(tree$Nnode+length(tree$tip))
  
  # adding possible offset if tree fully extinct
  if(!is.null(tree$root.time)) node.ages = node.ages + tree$root.time - max(node.ages)
  
  return(node.ages)
}
```
