## Supplementary material for "Ignoring stratigraphic age uncertainty leads to erroneous estimates of species divergence times under the fossilized birth-death process": mammals_analysis.R

```
# full analysis of BEAST2 output on simulated dataset
mammals_analysis = function(treefile, outfolder, resultfolder) {
  tmpfile = paste0(basename(outfolder),".RData")
  repeat {
    partial1 = mammals_allnodes_divtimes(treefile, outfolder, tmp_file = tmpfile)
    if(class(partial1) != "try-error") break
  } 
  partial2 = mammals_log_analysis(treefile, outfolder)
  partial = c(partial1, partial2)
  psummary = mammals_summary(partial)
  
  names = c("divtime", "diversificationRateFBD","turnoverFBD", "samplingProportionFBD", "SACountFBD")
  names = c(names, "rate.mean")
  full = fsummary = list()
  for (nm in names) {
    full[[nm]] = partial[which(grepl(nm, names(partial)))]
    fsummary[[nm]] = psummary[which(grepl(nm, names(psummary)))]
  }
  full$ESS = partial$ESS
  fsummary$ESS = psummary$ESS
  
  assign(paste0(basename(outfolder),"_accuracy"), full)
  save(list = c(paste0(basename(outfolder),"_accuracy")), file = paste0(resultfolder,basename(outfolder),"_accuracy.RData"))
  assign(paste0(basename(outfolder),"_summary"), fsummary)
  save(list = c(paste0(basename(outfolder),"_summary")), file = paste0(resultfolder,basename(outfolder),"_summary.RData"))
  file.remove(tmpfile)
  
  perf = mammals_performance(outfolder)
  sperf = lapply(perf, mean)
  assign(paste0(basename(outfolder),"_performance"),perf)
  assign(paste0(basename(outfolder),"_performance_summary"),sperf)
  save(list = c(paste0(basename(outfolder),"_performance"),paste0(basename(outfolder),"_performance_summary")), file = paste0(resultfolder,basename(outfolder),"_performance.RData"))
}
```
