## Supplementary material for "Ignoring stratigraphic age uncertainty leads to erroneous estimates of species divergence times under the fossilized birth-death process": mammals_performance.R

```
# analyze performance of runs 
# NB: based on Euler cluster, which records runtimes in LSF files
mammals_performance = function(outfolder, idx = 1:100, morph = F) {
  library(ape)
  library(readr)
  
  result = list()
  names = c("correct_ages", "median_ages", "interval_ages", "random_ages", "norm_intvl_ages")
  CPUtimes = list()
  for(nm in names) CPUtimes[[nm]] = rep(0, length(idx))
  
  # figure out CPU time from all lsf files
  lsfs = list.files(outfolder, pattern = "lsf", full.names = T)
  for(lsf in lsfs) {
    text = read_lines(lsf)
    
    time = grep("CPU time", text, value = T)
    time = strsplit(time," ")[[1]]
    time = as.numeric(time[length(time) - 1])
    
    filenm = if(!morph) grep("^/cluster/home/bjoelle/beast", text, value = T) else grep("^java -Xmx1024M -jar", text, value = T)
    filenm = strsplit(filenm," ")[[1]]
    filenm = strsplit(filenm[length(filenm)],"_")[[1]]
    
    i = as.integer(strsplit(filenm[length(filenm)], ".", fixed = T)[[1]][1])
    nm = paste0(filenm[c(-1,-length(filenm))],collapse = "_")
    
    CPUtimes[[nm]][i] = CPUtimes[[nm]][i] + time
    #print(paste(nm,i))
  }
  
  for(nm in names) result[[paste0(nm,".cpu_time")]] = CPUtimes[[nm]]
  
  for(i in idx) {
    for (nm in names) {
      print(paste(nm,i))
      # calculate ESS on tree total length
      log = read.table(file.path(outfolder, paste0("FBD_", nm,"_",i,".log")),header = T,comment.char = "#")
      log = log[(round(nrow(log)/10)+1):nrow(log),]
      ESS_h = ESScalc(log$TreeHeight)
      ESS_p = ESScalc(log$posterior)
      
      #save results
      result[[paste0(nm,".ess_hg")]] = c(result[[paste0(nm,".ess_hg")]], ESS_h)
      result[[paste0(nm,".perf_hg")]] = c(result[[paste0(nm,".perf_hg")]], CPUtimes[[nm]][i]/ESS_h)
      result[[paste0(nm,".ess_pos")]] = c(result[[paste0(nm,".ess_pos")]], ESS_p)
      result[[paste0(nm,".perf_pos")]] = c(result[[paste0(nm,".perf_pos")]], CPUtimes[[nm]][i]/ESS_p)
    }
  }
  result
}
```
