## Supplementary material for "Ignoring stratigraphic age uncertainty leads to erroneous estimates of species divergence times under the fossilized birth-death process": README.R

source("mammals_analysis.R")
source("mammals_performance.R")
source("mammals_accuracy.R")

# To do the full analysis of BEAST2 output on a simulated dataset
treefile = "path/to/dataset/stored/as/RData"
outfolder = "path/to/BEAST2/logfiles"
resultfolder = "path/to/store/results"
mammals_analysis(treefile, outfolder, resultfolder)
