## Supplementary material for "Ignoring stratigraphic age uncertainty leads to erroneous estimates of species divergence times under the fossilized birth-death process": figures.R

```
cetaceans_plot = function(outfolder, plotfile = NULL) {
  library(ggplot2)
  estnames = c("diversificationRateFBD","turnoverFBD", "samplingProportionFBD")
  pnames = c("Diversification rate","Turnover", "Sampling proportion")
  names = c("median_ages", "random_ages", "interval_ages")
  
  logs = list()
  for(x in names) {
    logs[[x]] = read.table(file.path(outfolder, paste0("Cetaceans_", x,".log")),header = T,comment.char = "#")
    n = nrow(logs[[x]])
    logs[[x]] = logs[[x]][(round(n/10)+1):n,]
  }
  
  p = list()
  for(i in 1:length(estnames)) {
    e = estnames[i]
    df = data.frame(values = c(), ds = c())
    for(x in names) {
      df = rbind(df, data.frame(values = logs[[x]][[e]], ds = x))
    }
    px = ggplot(df, aes(x = ds, y = values, fill = ds)) + geom_boxplot(notch = TRUE) + theme_bw() +
      scale_x_discrete(labels = c("Median ages", "Random ages", "Interval ages")) + xlab("") + ylab("") + 
      ggtitle(pnames[i]) + theme(legend.position = "none") + theme(axis.text.x = element_text(angle = 45, hjust = 1))
    p[[i]] = px
  }
  
  p = gridExtra::grid.arrange(grobs = p, ncol = 3)
  if(!is.null(plotfile)) ggsave(plotfile, plot = p,width = 10, height = 5)
  else print(p)
}

figure_ranges = function(file, plotfile = NULL) {
  load(file)
  data = get("pbdb")
  
  df = data.frame(id=1:length(data[,1]), min = data[,1], max = data[,2])
  library(ggplot2)
  pl = ggplot(df, aes(x=id))+ xlab("") + ylab("Time before the present (My)") +
    geom_linerange(aes(ymin=min,ymax=max),linetype=1,color="blue")+
    geom_point(aes(y=min),size=1,color="red")+
    geom_point(aes(y=max),size=1,color="red")+
    theme_bw()
  
  if(!is.null(plotfile)) ggsave(paste0(plotfile,"_ranges.pdf"))
  else print(pl)
}

plot_comparison = function(file, plotfolder, names = c("divtime", "diversificationRateFBD","turnoverFBD", "samplingProportionFBD")) {
  library(ggplot2)
  load(file)
  outname = tools::file_path_sans_ext(basename(file))
  fullfile = get(tools::file_path_sans_ext(basename(file)))
  
  plotnames = c("Divergence time", "Diversification rate","Turnover", "Sampling proportion")
  names(plotnames) = c("extmrca.divtime", "diversificationRateFBD","turnoverFBD", "samplingProportionFBD")
  
  for(name in names) {
    full = fullfile[[name]]
    cats = c("correct_ages", "median_ages", "interval_ages", "random_ages", "norm_intvl_ages")
    if(name == "divtime") name = "extmrca.divtime"
    df = data.frame(Age = c("Correct age", "Median age", "Interval ages", "Random age", "Symmetric ages"), 
                    Relative_error = sapply(cats, function(x) {full[[paste0(x,".",name,".relative_error")]]}),
                    Coverage = sapply(cats, function(x) {full[[paste0(x,".",name,".coverage")]]}))
    
    p2 <- ggplot(df, aes(x = Age, y = Relative_error)) + geom_point() + xlab(plotnames[name]) + #theme_bw() +
      theme(axis.text.x = element_text(angle = 45, vjust = 1, hjust=1), text = element_text(size = 15)) + ylim(0,0.8) + ylab("Relative error")
      
    ggsave(paste0(outfolder,outname, "_",name,"_rel_error.pdf"),width = 5,height = 5)
    
    p2 <- ggplot(df, aes(x = Age, y = Coverage)) + geom_point() + xlab(plotnames[name]) + #theme_bw() +
      theme(axis.text.x = element_text(angle = 45, vjust = 1, hjust=1), text = element_text(size = 15)) + ylim(0,1)
      
    ggsave(paste0(outfolder,outname, "_",name,"_coverage.pdf"),width = 5,height = 5)
  }
}

plot_node_comparison = function(log_int, log_med, log_rnd, plotfile = NULL, string = NULL) {
  library(ggplot2)
  if(!is.null(string)) {
    log_int = log_int[[string]]
    log_med = log_med[[string]]
    log_rnd = log_rnd[[string]]
  }
  
  df = data.frame(values = log_int, Age = "Interval ages")
  df = rbind(df, data.frame(values = log_med, Age = "Median ages"))
  df = rbind(df, data.frame(values = log_rnd, Age = "Random ages"))
  
  p = ggplot(df, aes(x = values, y=..scaled.., col = Age)) + geom_density(n = 2^8) + theme_bw() + xlab("") + ggtitle("Divergence time of Phocoenidae") +
    geom_hline(color = "black", yintercept = 0, size = 0.5) + xlim(12,22) + theme(plot.title = element_text(hjust = 0.5)) + ylab("Density (scaled to [0;1])")
  
  if(is.null(plotfile)) print(p)
  else ggsave(plotfile)
}
```
