## Supplementary material for "Ignoring stratigraphic age uncertainty leads to erroneous estimates of species divergence times under the fossilized birth-death process": README.R

source("figures.R")

# To make a plot of age ranges
outfile = "path/to/plot"
pbdbfile = "path/to/ranges/RData" #ranges should be stored as a table with minimum (1st col) and maximum (2nd col) age 
figure_ranges(pbdbfile, outfile)

# To make a plot comparing relative errors and coverage on simulated data
summary_file = "file/with/estimate/error/and/coverage" # as produced by mammals_summary
plotfolder = "path/to/figures/folder"
plot_comparison(summary_file, plotfolder)

# To make a plot comparing estimates of FBD parameters on cetaceans
outfolder = "path/to/folder/with/logfiles"
plotfile = "path/to/output/figure"
cetaceans_plot(outfolder, plotfile)

# To make a plot comparing posteriors on one node
log_int = "vector/containing/node/height/with/interval/ages"
log_med = "vector/containing/node/height/with/median/ages"
log_rnd = "vector/containing/node/height/with/random/ages"
plotfile = "path/to/output/figure"
plot_node_comparison(log_int, log_med, log_rnd, plotfile)
