## Supplementary material for "Ignoring stratigraphic age uncertainty leads to erroneous estimates of species divergence times under the fossilized birth-death process": auxf.R

```
sim.sequences = function(tree) {
  seqs = list()
  ntips = length(tree$tip.label)
  internal_branches = (ntips*2) - 2
  
  #for(j in 1:loci){
  # sample a mean substitution rate for each locus
  # under the gamma distribution
  mean.rate = rgamma(1, shape = 2, scale = 1/2)
  
  # simulate variable rates under a log normal distribution
  lstdev = 0.1
  simulated.rates = rlnorm(internal_branches, log(mean.rate), lstdev)
  
  # generate new branch lengths
  new.blens = tree$edge.length * simulated.rates /100
  
  # assign them to the tree
  new.tree = tree
  new.tree$edge.length = new.blens
  
  seqs = phyclust::seqgen(opts = paste(model, " -l", alignment.length, sep = ""), newick.tree = ape::write.tree(new.tree)) [-1]
  #}
  for(i in 1:length(seqs)) {
    tmp = strsplit(seqs[i],split = " ")[[1]]
    names(seqs)[i] = tmp[1]
    seqs[i] = tmp[length(tmp)]
  }
  return(list(seqs = seqs, mean.rate = mean.rate/100))
}

get_intervals_from_PBDB = function(f) {
  load(pbdbf)
  max_ages = c()
  min_ages = c()
  for (i in 1:length(f$h)) {
    t = f$h[i]
    idxes = intersect(which(pbdb$min.age <= t), which(pbdb$max.age >= t))
    
    if(length(idxes) == 0) {
      tp = runif(1,0,mean_pbdb_span)
      max_ages[i] = t + tp
      min_ages[i] = max(t - mean_pbdb_span + tp, 0)
    }
    else {
      if(length(idxes) > 1) {
        idxes = sample(idxes, 1, prob = pbdb$count[idxes]/sum(pbdb$count[idxes]))
      }
      max_ages[i] = pbdb$max.age[idxes]
      min_ages[i] = pbdb$min.age[idxes]
    }
  }
  
  list(max = max_ages, min = min_ages)
}

.find_clades = function(tree, nextant) {
  clades = list()
  aux = function(node) {
    desc = which(tree$edge[,1] == node)
    if(length(desc) == 0) return(node)
    
    c1 = aux(tree$edge[desc[1],2])
    c2 = aux(tree$edge[desc[2],2])
    
    c1.hasf = any(c1 > nextant)
    c2.hasf = any(c2 > nextant)
    c1.hase = any(c1 <= nextant)
    c2.hase = any(c2 <= nextant)
    
    if( c1.hase && c1.hasf && c2.hase ) clades[[length(clades) + 1]] <<- c1
    if( c2.hase && c2.hasf && c1.hase ) clades[[length(clades) + 1]] <<- c2
    
    c(c1,c2)
  }
  
  aux(length(tree$tip.label) + 1)
  clades
}

### make a rate matrix
rate_vect <- function(nchars, alpha){
  rbeta(nchars, alpha, alpha)
}

### create complete character matrix - all characters assumed symmetric
sim_character_sequences = function(tree, nchars, prop_binary, rate = NULL, alpha = NULL) {
  if(!is.null(alpha)) { 
    rates = rate_vect(nchars, alpha)
  } else {
    rates = rep(rate,nchars)
  }
  mats = lapply(rates, function(x) {
    if(runif(1) < prop_binary) rbind(c(-x, x), c(x, -x)) #binary character
    else rbind(c(-x, x/2, x/2), c(x/2, -x, x/2), c(x/2, x/2, -x)) #trinary character
  })
  seq = geiger::sim.char(tree, mats, model = "discrete")[,,1]
  seq = apply(seq, 1, function(x) paste0(x - 1, collapse = ""))
  seq
}
```
