## Supplementary material for "Ignoring stratigraphic age uncertainty leads to erroneous estimates of species divergence times under the fossilized birth-death process": mammals_simulation.R

```
### Simulation parameters

spec_rate = 0.15
ext_rate = 0.1
min_age = 40
max_age = 100

ntrees = 100
nextant = 25

sampl_rate = 0.2
min_fossils = 4
max_fossils = 125

alignment.length = 2000
loci = 1
model = "-mHKY -t5 -a0.25 -g5"

xml_template = "mammals_FBD_template.xml" # xml template should contain all components except alignment, ages & topological constraints
out_dir = "path/to/folder/for/xmls"
save.file = "path/to/save/dataset/as/RData"
pbdbf = "path/to/ranges/as/RData" # ranges should be a table with minimum age (1st col), maximum age (2nd col) and count (3rd col)

mean_pbdb_span = 8
set.seed(451)

### End of simulation parameters

source("auxf.R")
if(!dir.exists(out_dir)) dir.create(out_dir)

# simulating trees and fossils with parameters
trees = list()
fossils = list()
samp_trees = list()
while(length(trees) < ntrees) {
  nsim = ntrees - length(trees)
  trees = c(trees, TreeSim::sim.bd.taxa(nextant, nsim, spec_rate, ext_rate,complete = T))
  for (i in ntrees:(ntrees-nsim+1)) {
    origin = max(ape::node.depth.edgelength(trees[[i]])) + trees[[i]]$root.edge
    if(origin > max_age || origin < min_age) {
      trees = trees[-i]
      fossils = fossils[-i]
      next
    }
    
    fossils[[i]] = FossilSim::sim.fossils.poisson(sampl_rate, tree = trees[[i]])
    if(length(fossils[[i]]$edge) < min_fossils || length(fossils[[i]]$edge) > max_fossils) {
      fossils = fossils[-i]
      trees = trees[-i]
    }
  }
}

library(xml2)
sequences = list()
rates = c()
for (i in 1:ntrees) {
  # simulating sequences on the trees
  tmp = sim.sequences(trees[[i]])
  sequences[[i]] = tmp$seqs
  rates[i] = tmp$mean.rate
  
  # adding uncertainty to fossil ages
  fossils[[i]]$h = (fossils[[i]]$hmin + fossils[[i]]$hmax)/2
  fossils[[i]] = fossils[[i]][order(fossils[[i]]$edge, -fossils[[i]]$h),]
  intervals = get_intervals_from_PBDB(fossils[[i]])
  max.ages = intervals$max
  min.ages = intervals$min
  
  ftree = combined.tree.with.fossils(trees[[i]],fossils[[i]])
  tree = sampled.tree.from.combined(ftree)
  samp_trees[[i]] = tree
  ages = ape::node.depth.edgelength(tree)
  ages = max(ages) - ages
  
  # create xml for certain and uncertain runs
  template = read_xml(xml_template)
  dta = xml_find_first(template,"data")
  tax_ref = rep(F,length(tree$tip.label))
  text = ""
  text_med = ""
  text_rand = ""
  
  # handling sequences & tip dates
  for(j in 1:length(tree$tip.label)) {
    tip_id = as.character(tree$tip.label[j])
    name = strsplit(tip_id,split = "_")[[1]][1]
    if(j <= nextant) xml_add_child(dta, as_xml_document(list(sequence = structure(list(), id = paste0("seq_", tip_id), taxon = tip_id, totalcount="4", 
                                                                                  value = sequences[[i]][name]))))
    else xml_add_child(dta, as_xml_document(list(sequence = structure(list(), id = paste0("seq_", tip_id), taxon = tip_id, totalcount="4", 
                                                                      value = paste0(rep("?", alignment.length), collapse = "")))))
    if(j > 1) {
      text = paste0(text, ",")
      text_med = paste0(text_med, ",")
      text_rand = paste0(text_rand, ",")
    }
    text = paste0(text, "\n", tip_id, "=", round(ages[j],9))
    if(j > nextant) {
      text_med = paste0(text_med, "\n", tip_id, "=", (max.ages[j-nextant]+min.ages[j-nextant])/2)
      text_rand = paste0(text_rand, "\n", tip_id, "=", runif(1,min.ages[j-nextant],max.ages[j-nextant]))
    }
    else {
      text_med = paste0(text_med, "\n", tip_id, "=", 0)
      text_rand = paste0(text_rand, "\n", tip_id, "=", 0)
    }
  }
  xml_replace(xml_find_first(template,"data"),dta)
  
  # MRCA priors on fossils
  clades = .find_clades(tree, nextant)
  log = xml_find_first(template,"run/logger")
  dist = xml_find_first(template,"run/distribution/distribution[@id='prior']")
  for(x in clades) {
    child = as_xml_document(list(distribution = structure(list(),
      id = paste0(paste0(x,collapse = "."),".prior"), spec="beast.math.distributions.MRCAPrior", monophyletic="true", tree="@Tree.t:26")))
    taxset = as_xml_document(list(taxonset = structure(list(), id = paste0(x,collapse = "."), spec="TaxonSet")))
    for (tip in x) {
      if(!tax_ref[tip]) xml_add_child(taxset, as_xml_document(list(taxon = structure(list(), id=tree$tip.label[tip], spec="Taxon"))))
      else xml_add_child(taxset, as_xml_document(list(taxon = structure(list(), idref=tree$tip.label[tip]))))
    }
    xml_add_child(child,taxset)
    xml_add_child(dist, child)
    tax_ref[x] = T
    
    xml_add_child(log, as_xml_document(list(log = structure(list(), idref = paste0(paste0(x,collapse = "."),".prior")))))
  }
  xml_replace(xml_find_first(template,"run/distribution/distribution[@id='prior']"), dist)
  xml_replace(xml_find_first(template,"run/logger"),log)
  
  # different tip dates & names for median & correct ages
  trait = xml_find_first(template,"run/state/tree/trait")
  xml_text(trait) = text
  xml_replace(xml_find_first(template,"run/state/tree/trait"), trait)
  
  xml_attr(log, "fileName") = paste0("FBD_correct_ages_",i,".log")
  xml_replace(xml_find_first(template,"run/logger"),log)
  treelog = xml_find_first(template, "run/logger[@mode='tree']")
  xml_attr(treelog, "fileName") = paste0("FBD_correct_ages_",i,".trees")
  xml_replace(xml_find_first(template, "run/logger[@mode='tree']"),treelog)
  write_xml(template,file.path(out_dir,paste0("FBD_correct_ages_",i,".xml")))
  
  template2 = template
  xml_text(trait) = text_med
  xml_replace(xml_find_first(template2,"run/state/tree/trait"), trait)
  
  xml_attr(log, "fileName") = paste0("FBD_median_ages_",i,".log")
  xml_replace(xml_find_first(template2,"run/logger"),log)
  xml_attr(treelog, "fileName") = paste0("FBD_median_ages_",i,".trees")
  xml_replace(xml_find_first(template2, "run/logger[@mode='tree']"),treelog)
  write_xml(template2,file.path(out_dir,paste0("FBD_median_ages_",i,".xml")))
  
  template3 = template
  xml_text(trait) = text_rand
  xml_replace(xml_find_first(template3,"run/state/tree/trait"), trait)
  
  xml_attr(log, "fileName") = paste0("FBD_random_ages_",i,".log")
  xml_replace(xml_find_first(template3,"run/logger"),log)
  xml_attr(treelog, "fileName") = paste0("FBD_random_ages_",i,".trees")
  xml_replace(xml_find_first(template3, "run/logger[@mode='tree']"),treelog)
  write_xml(template3,file.path(out_dir,paste0("FBD_random_ages_",i,".xml")))
  
  # adding uncertain ages
  operator = as_xml_document(list(operator = structure(list(), spec="SampledNodeDateRandomWalker", 
                                                       windowSize="10.", tree="@Tree.t:26", weight="5.")))
  operator_n = as_xml_document(list(operator = structure(list(), spec="SampledNodeDateRandomWalker", 
                                                       windowSize="10.", tree="@Tree.t:26", weight="5.")))
  taxset = as_xml_document(list(taxonset = structure(list(), id = "fossilTaxa", spec="TaxonSet")))
  for(j in (nextant+1):length(tree$tip.label)) {
    if(ages[j] < 1e-5) stop("Ordering problem")
    
    tip_id = as.character(tree$tip.label[j])
    if(!tax_ref[j]) xml_add_child(taxset, as_xml_document(list(taxon = structure(list(), id=tip_id, spec="Taxon"))))
    else xml_add_child(taxset, as_xml_document(list(taxon = structure(list(), idref=tip_id))))
    
    xml_add_child(operator, as_xml_document(list(samplingDates = structure(list(),
      id = paste0("samplingDate.",tip_id), spec="beast.evolution.tree.SamplingDate", taxon = paste0("@",tip_id),
      lower=as.character(min.ages[j - nextant]), upper=as.character(max.ages[j - nextant])))))
    xml_add_child(operator_n, as_xml_document(list(samplingDates = structure(list(),
      id = paste0("samplingDate.",tip_id), spec="beast.evolution.tree.SamplingDate", taxon = paste0("@",tip_id),
      lower=as.character(max(0, ages[j] - mean_pbdb_span/2)), upper=as.character(ages[j] + min(ages[j],mean_pbdb_span/2))))))
  }
  
  xml_add_child(operator, taxset)
  run = xml_find_first(template,"run")
  xml_add_child(run,operator)
  xml_replace(xml_find_first(template,"run"), run)
  
  xml_attr(log, "fileName") = paste0("FBD_interval_ages_",i,".log")
  xml_replace(xml_find_first(template,"run/logger"),log)
  xml_attr(treelog, "fileName") = paste0("FBD_interval_ages_",i,".trees")
  xml_replace(xml_find_first(template, "run/logger[@mode='tree']"),treelog)
  write_xml(template,file.path(out_dir,paste0("FBD_interval_ages_",i,".xml")))
  
  xml_add_child(operator_n, taxset)
  xml_replace(xml_find_first(run,"operator[@spec='SampledNodeDateRandomWalker']"),operator_n)
  xml_replace(xml_find_first(template,"run"), run)
  
  xml_attr(log, "fileName") = paste0("FBD_norm_intvl_ages_",i,".log")
  xml_replace(xml_find_first(template,"run/logger"),log)
  xml_attr(treelog, "fileName") = paste0("FBD_norm_intvl_ages_",i,".trees")
  xml_replace(xml_find_first(template, "run/logger[@mode='tree']"),treelog)
  write_xml(template,file.path(out_dir,paste0("FBD_norm_intvl_ages_",i,".xml")))
}

save(trees, fossils, samp_trees, rates, sequences, file = save.file)
```
