## Supplementary material for "Ignoring stratigraphic age uncertainty leads to erroneous estimates of species divergence times under the fossilized birth-death process": README.txt

This folder contains the data and code used in our article.

It is divided in the following subfolders:
- cetaceans_data contains the data related to the empirical cetaceans dataset
- code_files contains the R scripts used to simulate datasets, process the results and build the plots
- simulated_data contains the simulated datasets and results
