## Supplementary material for "Ignoring stratigraphic age uncertainty leads to erroneous estimates of species divergence times under the fossilized birth-death process": priors_mammals_beast2.pdf

### List of priors used for the simulated dataset.

| Parameter | Prior distribution |
|---|---|
| FBD diversification rate | Uniform$[0; \infty[$ |
| FBD turnover | Uniform$[0; 1[$ |
| FBD sampling proportion | Uniform$[0; 1[$ |
| Origin of the FBD process | Uniform$[0; \infty[$ |
| HKY+$\Gamma$ $\kappa$ parameter | LogNormal(1.0,1.25) |
| HKY+$\Gamma$ shape of the $\Gamma$ distribution | Exponential(1.0) |
| Relaxed clock mean rate | Uniform$[0; \infty[$ |
| Relaxed clock standard deviation | Gamma($\alpha = 0.5396$,$\beta = 0.3819$) |
