## Supplementary material for "Ignoring stratigraphic age uncertainty leads to erroneous estimates of species divergence times under the fossilized birth-death process": README.txt

This folder contains the data elated to the simulated mammals dataset.

It contains the following files:
- mammals_seed451_dataset.RData contains the simulated trees, sequences and fossil occurrences
- mammals_seed451_accuracy.RData and mammals_seed451_performance.RData contain summaries of the results on respectively accuracy and performance on the simulated dataset.
- pbdb_data_NOA_mammals.csv contains the PDBD readout used to simulate realistic fossil age ranges
- priors_mammals_beast2.pdf contains a list of the priors used for running BEAST2 on this dataset
